# The fingering patterns in the epithelial layer control the gap closure rate via curvature-mediated force

**DOI:** 10.1101/2023.03.06.531411

**Authors:** Hyuntae Jeong, Jinwook Yeo, Seunghwa Ryu, Jennifer Hyunjong Shin

## Abstract

Closing gaps in cellular monolayers is a fundamental aspect of both morphogenesis and wound healing. This closure can be achieved through leader cell crawling or actomyosin-based contraction, depending on the size of the gap. Here, we focus on wounds whose closure is driven by interfacial instabilities, featuring both leader cell-driven fingers and actin-mediated contraction. Our proposed model predicts a positive correlation between the frequency of fingering and the overall speed of boundary closure. This fingering frequency is precisely regulated through the orchestration of cell density-driven pressure, cell-cell repulsions, and the initial curvature of the wound boundary. Our findings demonstrate an inverse correlation between fingering frequency and boundary curvatures, indicating a “self-control” mechanism for closure rates independent of the initial curvatures of the wound periphery. Notably, changes in curvature caused by fingering formation generate force that aids in the healing process.

## Introduction

Gap closure is a ubiquitous physiological phenomenon that occurs in response to external injuries or internal apoptotic events during morphogenesis and tissue homeostasis^1–5^. Proper re-epithelization is critical to close wounds, involving two modes of cellular migration: purse-string contraction of the wound edge and crawling of the boundary cells ^6,7^ Purse-string contraction involves actomyosin cable contraction that spans over multiple cells along the boundary, whereas leader cell crawling is achieved through the active emergence of lamellipodia^4,8–12^. Previous studies have demonstrated the complementary actions of these two modes during wound closure^10,13–17^. Smaller wounds or voids created by defects in the extracellular matrix (ECM) are closed predominantly by purse-string-like contractions ^15,16^. Wounds are considered small when 10~20 cells are aligned along the perimeter without any discontinuity in actomyosin cables. Larger wounds, on the other hand, are closed through the active crawling of leader cells at the boundary^10,14^.

Two distinct closing modes can, however, occur simultaneously in a cooperative manner to expedite the closure^18–21^. The synergistic effects from the coexisting two modes have been demonstrated both experimentally and computationally for small wounds^22–24^. *Ravasio et al*. demonstrated the contribution of crawling forces during purse-string contractions in the small wound by verifying a mathematical superposition of purse-string and crawling^22^. Furthermore, the *in silico* model by Staddon et al. confirmed the increase in closure speed when two modes contributed mutually, whose relative dominance depended on the size of local curvatures^23^. Both studies suggested the cooperative effects of two modes for small wounds of high enough curvatures (0.1~0.6μm^-1^) spanning over a short perimeter of 10~20 cells. On the other hand, larger wounds involve complex features like abrupt crawling protrusions of finger-like shape at the regions of discontinued actomyosin cables^25–27^. As the convex fingers extrude further, concave suspending-bridge-like actomyosin cables appear between the fingers, which resemble negative curvature bridges on small wound boundaries and are expected to contribute to overall wound closure through purse-string mechanisms as the concave strains developed^10,14,28−30^. Although the development of purse-string contractions following the outgrowth of fingers would be an important process for the closing of large wounds, the descriptions for fingering extrusions and contraction of bridges mostly remain in stochastic models^28,31,32^. However, *Vishwakarma M. et al*. recently discovered that the adjacent fingers were equally space along the boundary, whose length scale was similar to the correlation length of cellular forces within the monolayer^33^. These findings led us to postulate that the fingering extrusion must be governed by the physical forces of constituent cells in the layer as being a well-regulated phenomenon to close the gap. Here, we aim to elucidate this intriguing event based on a mathematical model and quantification tools for cellular dynamics.

In this study, we focus on the closing event of reasonably large wounds, typically of 100s-1000s μm in radius, where both modes of closure are important. To elucidate the closing mechanism, we first identify the role of fingers in wound closures based on a simple mathematical isotropic line tension model that reflects forces along the fingering patterns. Comparing the mathematical model and experimental results clarifies that fingering extrusion controls overall wound closure and is orchestrated by density gradient-driven cell flux and the initial boundary curvatures of the wound boundary.

## Results

### Initial diameter-dependent fingering protrusion dictates dynamic changes in wound boundary

In this study, the wounds are created by culturing the MDCK (Madin-Darby Canine Kidney cells) monolayer with a PDMS stencil for 18 hrs and carefully removing the stencils to obtain a smooth wound edge of the desired shape and size (Fig. 1(a)). We use circular wounds of various initial radii (*R_initial_*) ranging from 150μm to 500μm and a straight one to investigate the effect of the initial wound curvature (*κ_initial_* = 1/*R_initial_*) to confirm the previously reported correlation between the initial curvature and the wound closing modes^22^. Consistently with existing reports, the small wound (*R_initial_* = 150μm) shows a shrinking with a smooth boundary, predominantly via purse-string-like contraction. In contrast, the wound of larger radii (*R_initial_* =250, 500μm, ∞) shows a rough interface due to the actively protruding cells at the onset of the closure (Fig. 1(b)). As a reference, a 150μm radius (942μm in perimeter) corresponds to roughly ~30 cells, and a 500μm radius (3142μm in perimeter) includes ~100 cells. As shown in the boundary trajectories of a large wound (*R_initial_* = 500μm), the initially smooth boundary begins to exhibit periodic waves formed by the emergence of extruding fingers (Fig. 1(c)). As the fingers extruded further, a local concave bridge between two adjacent fingers is naturally developed, and this local concave region is termed “valley” henceforth. The difference in the advancing speeds of the closing boundary between the fast-moving fingers and the slow-moving valleys causes an increase in negative curvature of the valley (Fig. 1(d)). Soon after, valley regions begin to accelerate, possibly to avoid the excess negative curvatures, gradually smoothening the boundary trajectories as shown clearly in Fig. 1(d) from a dashed blue (t=7hr) line to a dotted green (t=8hr) line. Here, we define the roughness index (*RI*) of the wound boundary as the ratio between the contour perimeter and the convex hull perimeter (Fig. 1(e)). For the smallest wound closure (R=75μm), the *RI* persistently decreases throughout the closure, indicating a smooth shrinking over time without any finger formation. For larger wounds of *R_initial_* >150μm, however, the *RI* curves exhibit positive slopes during earlier times (0~6 hrs), reflecting the initial formation of fingers. Once the *RI* reaches its peak value, it slowly decays over time, which suggests the temporal maintenance of fingers and valleys (Fig. 1(e)). The maximum *RI* value and the duration over which higher *RI* value is maintained positively correlate with the *R_initial_*, which reflects the more prominent effect of the fingers on the closure of larger wounds. To investigate the basis for the *R_initial_*-dependent *RI* profile, we further quantify two major characteristics of the fingers, namely the finger amplitudes and the finger-to-finger distance (*FFD*). The amplitude is defined as the length of fingers from the inscribed circle of boundaries after 6 hrs from the protrusion. The *FFD* is the distance between the adjacent fingers. As shown in Fig. S1, the finger amplitude shows insignificant dependence on the *R_initial_*, whereas the *FFD* exhibits a notable negative correlation with the *R_initial_* (Fig. 1(f)), suggesting that the occurrence of the fingers predominantly influences the *RI* profile than their amplitudes. When the *FFD* is plotted as a function of time for *R_initial_* = 500μm, the formations and falls of the *FFD*, reflecting the genesis and merger of the fingers, clearly show slow decays similar with the maintenance of RI in larger wounds, where the red dotted line represents the mean value of 17 samples (Fig. 1(g)). On the other hand, the *FFD* changes for *R_initial_* = 150μm showed drastic decrease corresponding to the RI changes in small wounds, where the red dotted line represents the man value of 8 samples (Fig. 1(g)). The corresponding decaying of *FFD* values in the plot supports the changes of RI reflects the fingering dynamics along the boundary.

**Fig. 1.**
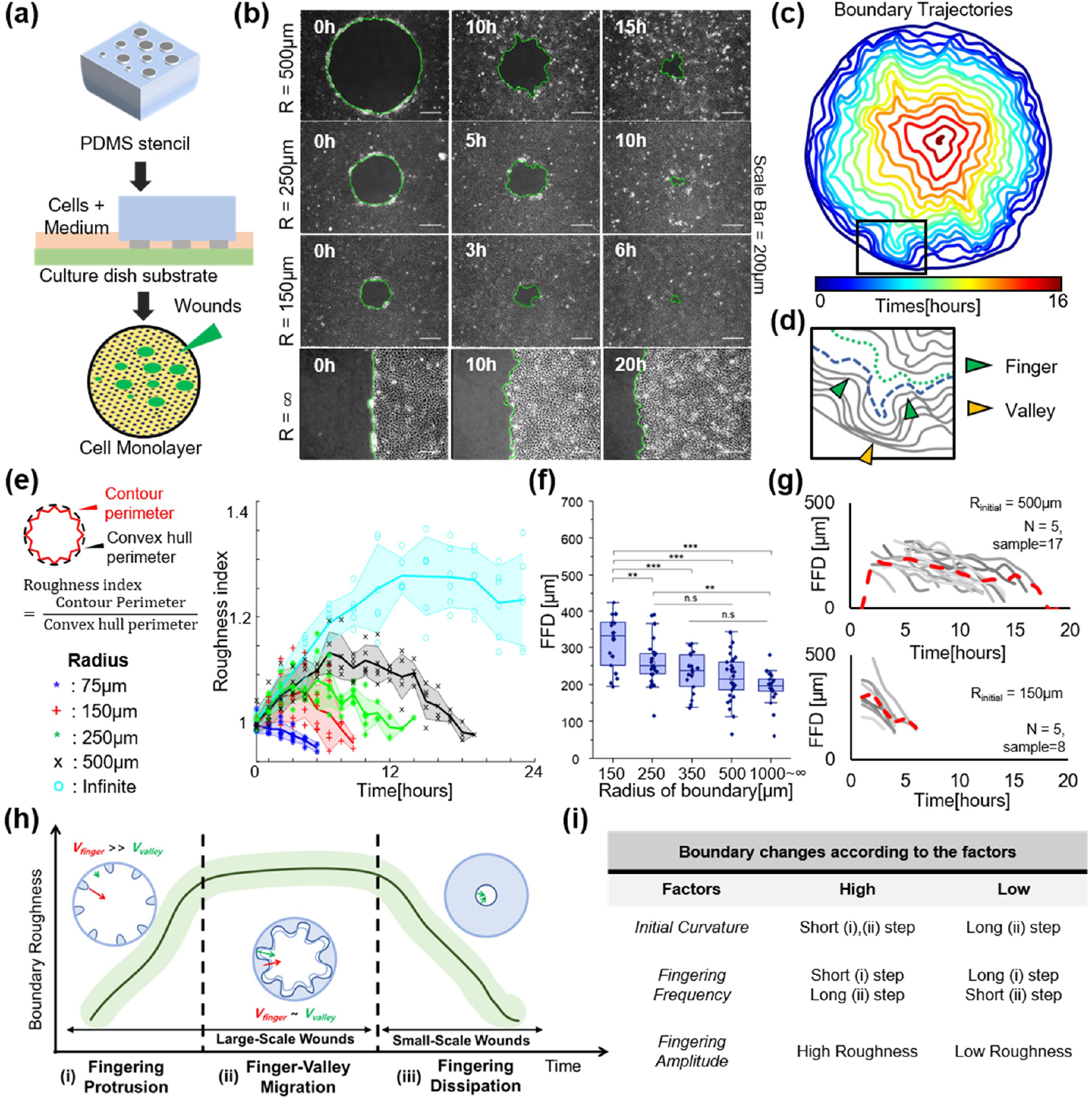
Fingering extrusions along the wound boundary with various curvatures formed by PDMS stencils. (a) Steps for creating various wound shapes by fabricating PDMS stencil from the silicon wafer. (b) Boundary shape changes during the wound closure according to initial diameters, smooth surface shrinking when the wound is comparatively small(radius = 150μm) and complicated change of wound boundary according to the fingering extrusion in the large wound(radius = 500μm, straight wound), (c) The trajectory of the boundary curves over time when the radius is 500μm, (d) Enlarged images of boundary trajectory when cells formed the fingering structures, green triangles indicate fingering regions and yellow triangles indicate valley regions, (e) Schematic for calculating the roughness of wound boundary and comparison of roughness of wound boundary with different radii (blue: 75μm, red: 150μm, green: 250 μm, black: 500μm, cyan: infinite), (f) Distance between fingers according to the initial diameters of the wounds (*: p ≤ 0.05, **: p≤**0.01**, ***: p ≤ 0.001), (g) Temporal change of distance between fingers during the wound closure (when the radius of wounds is 500μm). (h) Schematics for change of boundary shapes with three sequential steps due to the fingering extrusion, (i) Table for the changes of boundary shape according to the geometrical factors.

Based on these observations, we propose that wound closure in the epithelial cell layer occurs in three distinct regimes (Fig. 1(h)). During regime (i), the extrusion of fingers gradually increases the boundary roughness. Once wave-like sequential finger and valley structures develop, the rough boundary is maintained during regime (ii), as the valley regions follow the extruding fingertips. In the shrinking regime (iii), fingers disappear, and wound closure is accomplished by contraction of the smooth boundary. Especially, the initial boundary radius (*R_initial_*) and fingering characteristics, such as finger frequency (*FFD*) and finger amplitudes, dictate the closing process of wounds via controlling the span of each regime in the closure (Fig. 1(i)). Thus, the following sections of this paper extensively explore the roles of these variables on wound closure, with fingering extrusion serving as a key parameter for controlling the closing performance.

### A mathematical model for finger structures predicts the positive correlation between boundary speeds and fingering frequency

We first predict the role of finger-valley structures by simplifying the force along the boundary as an isotropic line tension (Fig. 2(a, b)), which counter-balances the protruding crawling force at the fingers. At the steady state where the boundary roughness is maintained at a constant level (regime (ii) of Fig. 1(h)), the force equilibrium is assumed between the crawling force at the fingers (*F*) and the line tension (*T*) at the boundary as follows,

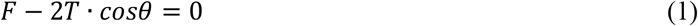

where *θ* is the angle between the vertical axis of fingertips and the tangent line of the finger boundary (Fig. 2(a)). By simplifying the valleys to spatially repetitive circular arcs, the force equilibrium of the valley region 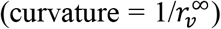 with a line tension and constant velocity (*ν*) of the boundary can be expressed as,

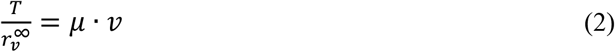

where the *μ* is a viscous friction coefficient, inclusive of resistance caused by both cell-cell and cell-substrate adhesions (Fig. 2(b)). To validate these relationships for given T and *μ*, we measure the velocity and boundary curvature 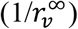 of the valley from the experimental data, confirming the linear correlation between these two quantities (Eq. 2) as shown in Fig. 2(c). Here, the instantaneous boundary velocity (*ν*) is measured by tracking the vertical displacement at a 1-hr interval, and the temporal changes in curvature are displayed by the color gradation in the plot. Interestingly, the finger velocities, marked by the purple-pink scale, are clustered around 30μm/h with no apparent dependency on the boundary curvature 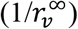 (Fig. 2(c)). These experimental data are well reflected on the spatiotemporal graph for the representative fingering extrusion that shows an almost constant slope as a function of time (Fig. 2(d)). In contrast, the valley region exhibits a distinct slope change at around 10hrs (Fig. 2(e)).

**Fig. 2.**
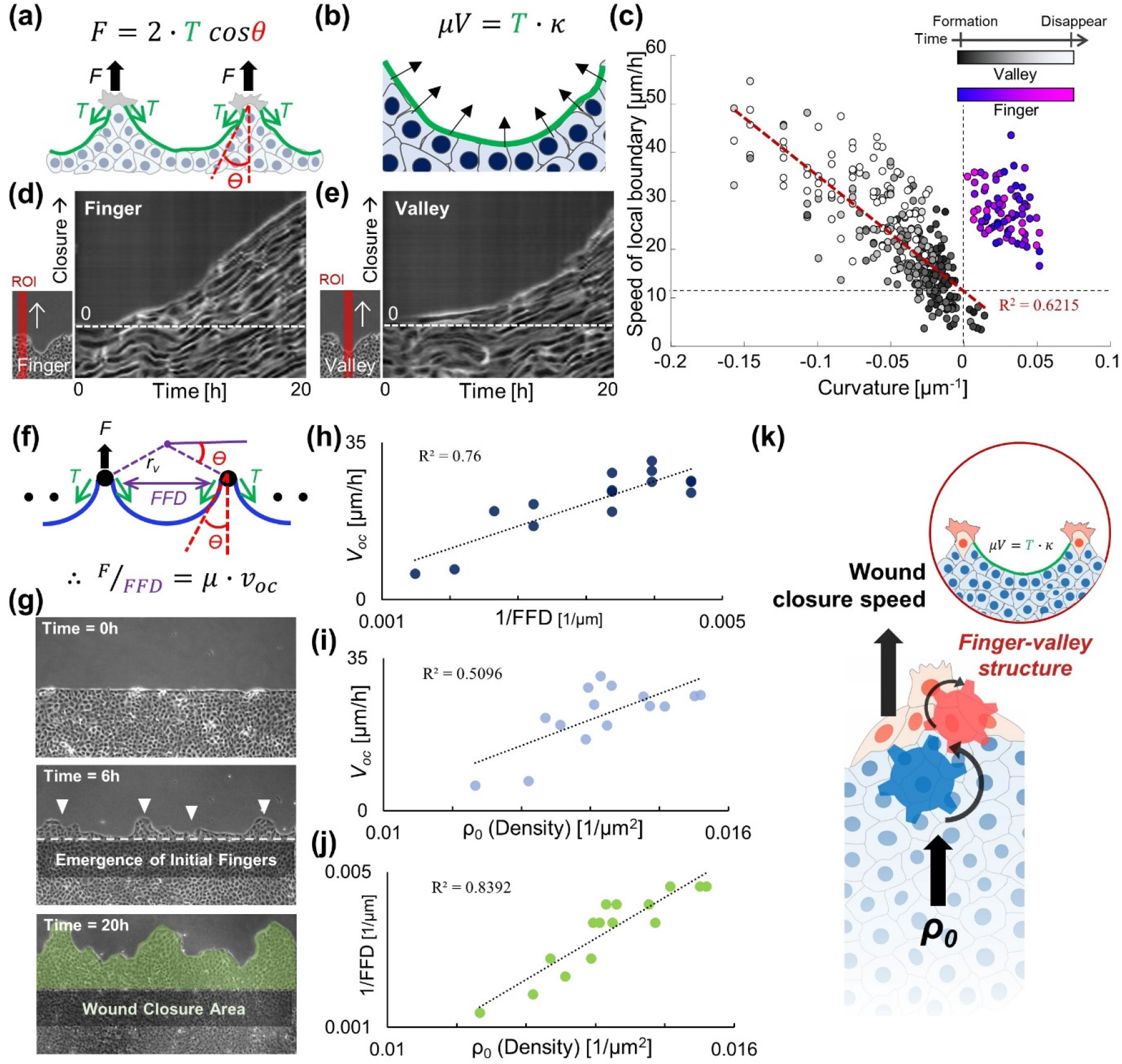
Effects of finger-valley structures on wound closure speeds based on the line tension model. (a, b) Schematics of force equilibrium at the finger-valley structures, (c) Distribution of proceeding speeds of wound edges according to the curvatures, the curvature was measured from the three points in the edge of valleys and fingers. The gray gradient and the color gradient indicate a relative time for diminishing of valleys and fingers, respectively. (d, e) Kymographs for proceeding of the finger and valley region of the straight-patterned wounds, (f) Schematics for the role of the fingering frequency on overall wound closure speeds, (g) Measurement of *ρ_0_, FFD* (at 6 hrs), and *V_oc_* (during 20hrs) for analyzing the correlation between variables, (h) Comparison of overall wound closure rate and the initial fingering frequency, (i) Comparison of overall wound closure rate and initial cell density in the monolayer (measured by counting the number of cells in windows), (j) Plot for a linear relationship between the initial density of cells and fingering frequency after 6 hrs, (k) Schematic for relationships amongst density-based flux, fingering extrusions, and the wound closure speed.

In the case of the overall wound closure, one can expect the repetitive extrusions of fingers to influence the overall closing speeds. From the relationship between the radius of curvature and the finger-to-finger distance (*FFD*), i.e., 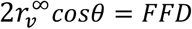, shown in Fig. 2(f), the boundary migration of straight wounds (*R_initial_* = ∞) can be expressed as the relationship between *F*, *FFD*, and the overall wound closure speeds *ν_oc_* as follows (Fig. 2(f)).

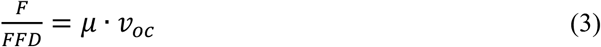

Here, Eq 3 lies on the assumption that tensional force *T* and friction coefficient *μ* are isotropic and constant with respect to local radius change, inferring the linear relationship between the *ν_oc_* to the fingering frequency (~1/*FFD*) in the straight wound geometry. To experimentally test the effect of initial fingering emergence on the overall wound closing rate, the average velocity values of the boundary measured over 20 hrs are plotted against 1/*FFD* of the corresponding samples at the moment where fingers begin to emerge (t = 6 hrs) as shown in Fig. 2 (g, h). The results confirm a clear positive correlation between the overall closing rate of the wound and the initial fingering frequency. Furthermore, given that cell density acts as a driving factor for bulk motion within cell monolayer, we investigate the effect of initial cell density (*ρo*) on the overall closing rate as shown in Fig. 2(i). Interestingly, the correlation strength between *νoc* and *ρo* is notably weaker than that between *ν_oc_* and 1/*FFD*, marked by the R^2^ value. On the other hand, *ρo* shows the strongest correlation (R^2^=0.8916) to the early fingering emergence (1/*FFD*) (Fig. 2(j)). The result implies that the *ρo* must act as an upstream cue that contributes to the emergence of the initial fingers, whose numbers eventually impact the overall *ν_oc_* as schematically illustrated in Fig. 2(k).

Thus far, we have clarified the direct role of fingers on the wound closure rate and identified the factors that regulate fingering frequency. Higher density is a primary candidate that increases the initial frequency of fingers (Fig. 2(j)), while the higher curvature of the boundary resists the emergence of fingers (Fig. 1(f)). Therefore, the following section will extensively analyze the role of these factors in the regulation of fingering frequency.

### Dynamic changes in cellular density induce cell fluxes for the fingering extrusions at the wound boundary

The cellular density (*ρ*), the first candidate known as a regulator for finger generation, is not homogeneous within the monolayer, both spatially and temporally (Fig. 3(a)). The spatial differences in densities create a local density gradient (∇*ρ*) within the monolayer that can perturb the velocity fields of cells due to the particle characteristics of diffusive migration. As shown in the series of cellular area (1/*ρ*) map in Fig. 3(b), the compressed cells in the high-density region develop the diverging velocity field towards the neighbors while their sizes relax to enlarge (Fig. 3(b)). The temporal plots for divergence and cellular area exhibit similar profiles, yet with a phase shift by ~2hrs, suggesting a possible role of the cellular density in developing the velocity divergence within the monolayer (Fig. 3(c)). To quantify the similarity between the two plots, we analyze the cross-correlation of divergence and unbiased cellular area calculated by subtracting the background decreasing trend caused by the increase in cell density. As shown in Fig. 3(d, e), the divergence and cellular area (1/*ρ*) demonstrate a fairly high correlation (*r_average_* = 0.55) with approximately 2 hrs time delay (*l_agaverage_* = 1h 45min). The cellular fluxes generated from the diverging source is transmitted to the boundary through intimate cell-cell junctions within the monolayer. This transmitted flux then can induce a thrusting pressure to form fingering extrusions at the boundary. To confirm this assumption, the spatiotemporal velocity changes in an ROI near the fingering extruding boundary in Fig. 3(f) are analyzed (Fig. 3(g-h)). The kymograph of velocity divergences exhibits several diagonal lines from the posterior region to the boundary, attributing to the spatiotemporal propagations of cell fluxes toward the boundary. Notably, the divergence source (red) is sequentially displaced toward the boundary until the emergence of the fingering extrusion (marked by red arrowheads). Consistently, diagonal lines in the kymograph of unbiased velocity obtained by removing mean velocities account for the propagation of fluxes from the inner monolayer leading to the formation of the fingers (Fig. 3(h)). Based on these quantified results, we confirm that the propagation of heterogeneous density-driven cell flux is the key mechanism for initiation of new finger extrusion at the boundary.

**Fig. 3.**
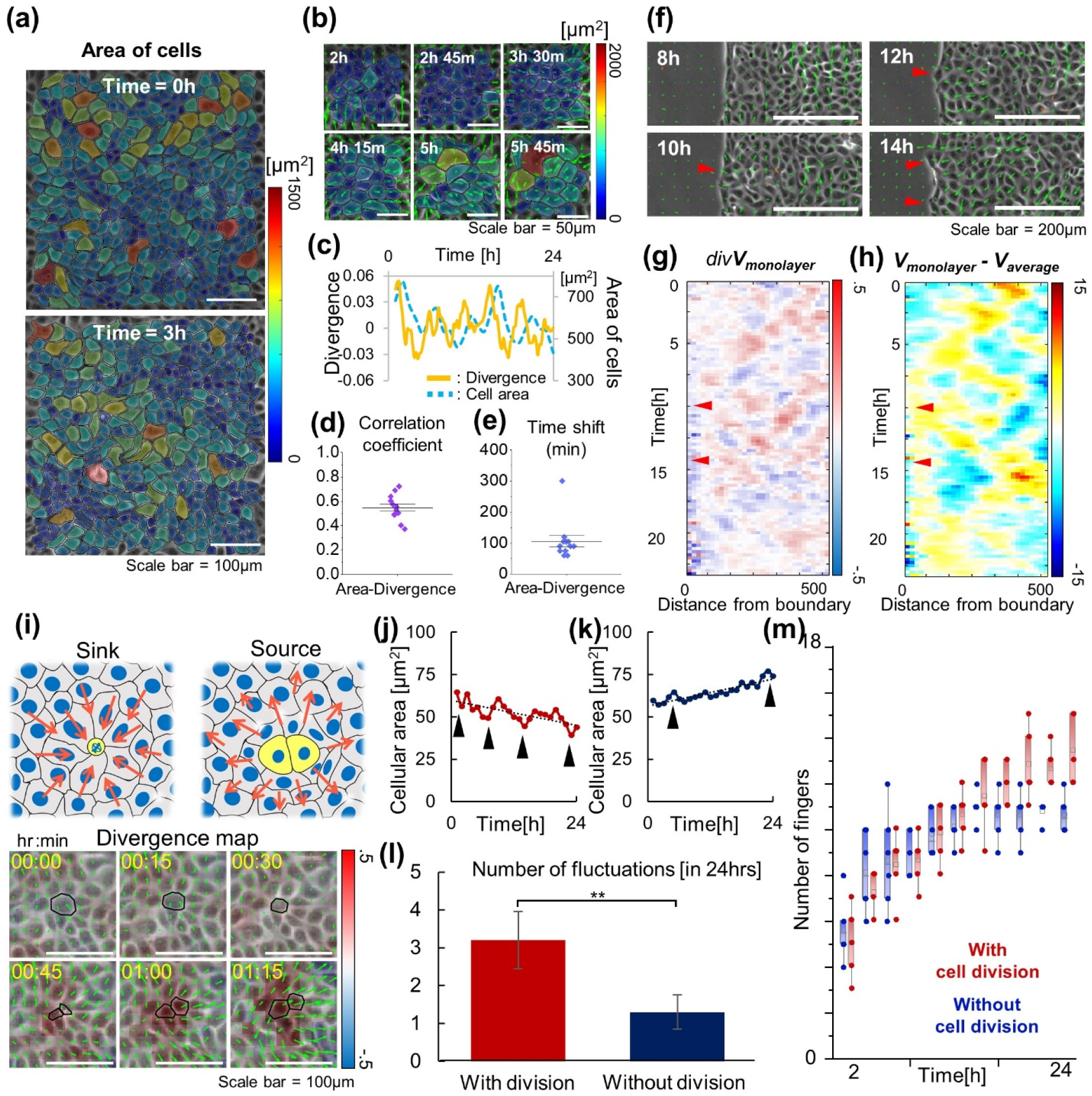
Dynamics of the cell populations in monolayer induced the fingering extrusions at the boundary. (a) Heterogeneously distributed cell density in the spatiotemporal domain, (b) Visualized results for the relationships between cellular density and cell fluxes (Green arrows = velocity vectors, Color map = Area of cells), (c) Comparison of changes in divergences and cellular area at the same domain during the wound closure, (d, e) Box plots for the correlation coefficient and time shift between divergence and cellular area plots, (f) Images of fingering extrusions in the region of interests, the red arrow heads indicate the location of fingering formations, (g, h) The kymograph of the divergence and unbiased velocity. The red arrows shows the time points when fingers are extruded. (i) Schematic for effects of proliferation for deriving disturbances of vector fields in the cell monolayer and divergence map with velocity vectors near the cell dividing region; the black line indicates the dividing cells. (j, k) Cellular area changes near the boundary (200×200μm2 from the edge), when the cells were dividing or not (Black arrow: each fluctuation). (l) Number of fluctuations when the cell division is controlled (*: p ≤ 0.05, **: p≤0.01, ***: p ≤ 0.001), (m) Changes in the number of fingers during the wound healing according to the cell proliferation conditions.

It is interesting to note that sporadically dividing cells naturally promote heterogeneity in densities, generating pressure against neighboring cells as the daughter cells expand (Fig. 3(i)). Given the highly proliferating nature of MDCK cells, it can be assumed that cell divisions sufficiently provoke divergences of velocity vectors, eventually leading to fingering extrusions. To test this hypothesis, we quantified the changes in cellular area (1/*ρ*) near the boundary while attenuating cell division through thymidine treatments. Thymidine arrests the cell cycle at the G1/S phase, thereby inhibiting cell division. As shown in Fig. 3(j-l), the mean cellular area shows periodic fluctuations in the untreated control conditions, whereas the mean cellular area continues to increase without fluctuations when cell division is inhibited by thymidine (Fig. S2). The emergence and propagation of velocity divergence (Fig. 3(g)) that act as cues for additional fingering extrusions also arise from dynamic fluctuations during the cell dividing process (Fig. S3, Mov. S1), resulting in an increase in the number of fingers only when cells are actively dividing (Fig. 3(m)). Inhibition of cell division naturally leads to the suppression of fingering formation in later time (>10hrs) (Fig. 3(m)). Conclusively, we have identified a critical role of cell division in the emergence of fingers by initiating divergences in the velocity vector field, acting as a local perturbation in density and velocity. Furthermore, in addition to the density-driven pressures, the initial polarity of traction forces also exhibits correlated results for the fingering positions. As shown in Fig. S4(a, b), the pre-polarized traction vectors to the normal direction to the boundary interface, which coincide with the crawling force of boundary cells, are shown to be localized at the immediate posterior region of the future fingers. This result suggests that the polarity of tractions near the boundary can determine where the crawling forces would accumulate to initiate the formation of fingers.

### Converging flux from the initially curved boundary does not increase the wound closure rate

Contrary to the straight wound boundary in previous sections, naturally formed wounds are likely to feature inherent initial curvatures (*κ_initial_*) that affect various aspects of wound closures tabulated in Fig. 1(i). As shown in Fig. 4(a), a negative *κ_initial_* at the boundary causes the converging flux of cells from the dense reservoir toward the center of the wound. In this scenario, the degree of convergence would increase with *κ_initial_*, which is predicted to result in a faster closure of the wounds (ΔA = constant, *ν_oc_* ∝ *κ_initial_*) with a higher fingering frequency. To test this prediction, we analyze the overall closure rate (*ν_oc_*) and ΔA as the parameters to assess the wound closure (Fig. 4(b)). The experimental data, however, show the positive linear relationship between ΔA and *R_initial_* (1/*κ_initial_*), contradicting our prediction (Fig. 4(c)). Consequently, the overall closure rate (ΔA/2π*R_initial_*) shows an almost independent relationship with *κ_initial_* (Fig. 4(d)). In addition, the previously obtained relationship between *FFD* and *R_initial_* (Fig. 1(f)) exhibits the opposite result to prediction. The number of cells between neighboring fingers, another parameter for the inverse of fingering frequency, also displays larger values as *κ_initial_* increases (Fig. S5). Since the fingering frequency has been found to linearly correlates with the wound closure speeds (Fig. 2 (f)), the *κ_initial_*-independent *ν_oc_* can be understood as a consequence of the counteractions between converging effects and downregulation of fingering extrusions at higher *κ_initial_* wound boundaries.

**Fig. 4.**
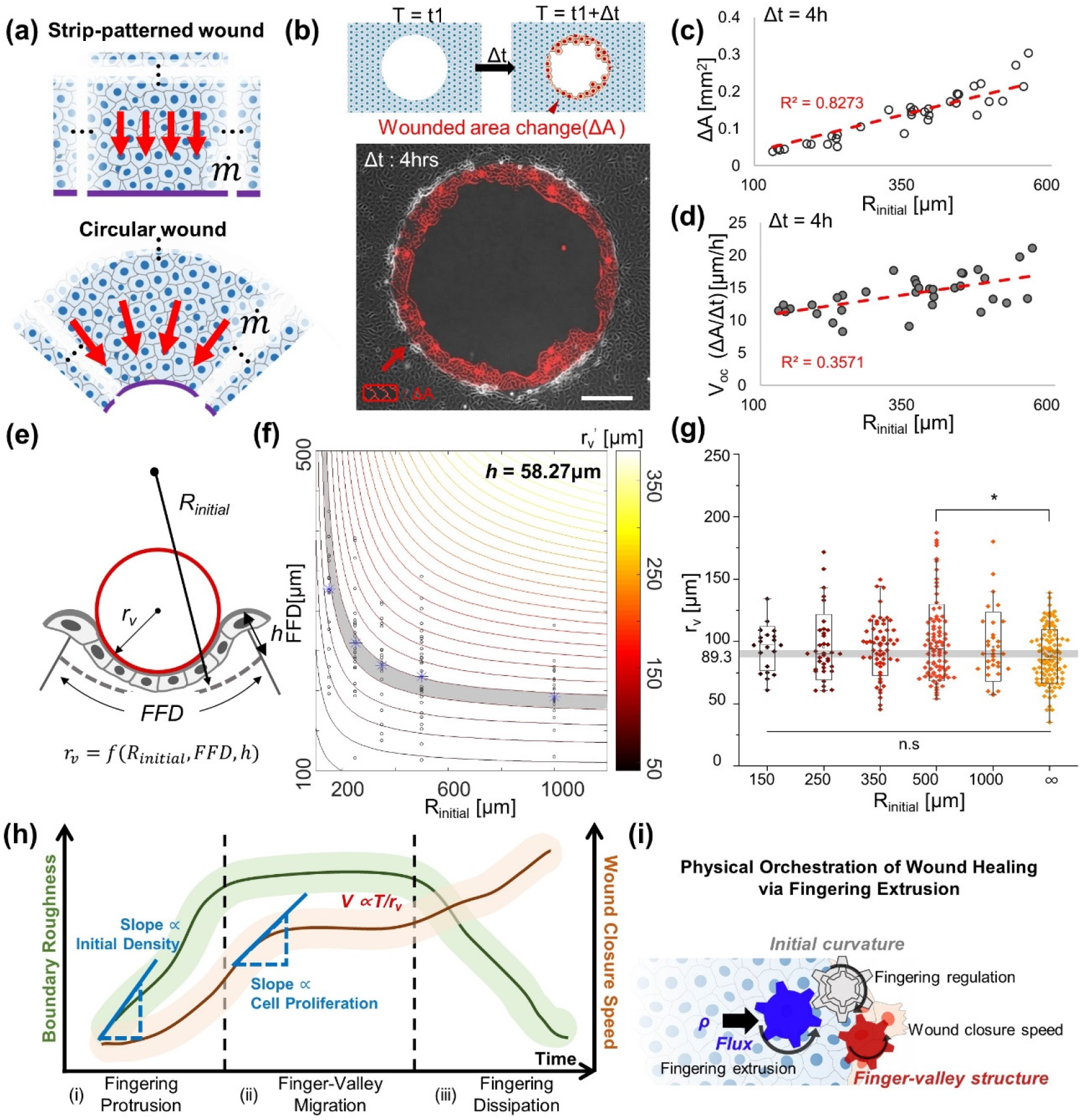
Changes in wound closure rates and fingering extrusions from the initial wound curvature differences. (a) Schematics for differences of the cell flux due to the initial curvature of boundary, (b) Measurement of wounded area changes in the experiments, the area between two boundaries with 4 hrs iteration is one data for wounded area changes. (c-d) Scatter plots for wounded area changes and mean wound healing speed according to the geometrical properties of the boundary, (e) Relationship amongst curvatures of valleys, FFD, initial curvature, and amplitudes of fingers(after fingers were formed), when cell boundary is initially curved, (f) Simulation results for relationships amongst final curvatures, initial diameters, and distance between finger, when *h* = 58.27 μm. And scatter plots of experiment results for FFD according to the R_initial_ (Blue stars: average value for each R_initial_), (g) Results of the local radius of curvature according to the initial boundary diameter. (*: p ≤ 0.05, **: p≤0.01, ***: p ≤ 0.001, determined by one-way ANOVA), (h) Schematic for the wound closing steps with regulating factors for fingering extrusion and developing speed, (i) Summary of wound closing mechanisms through the orchestrations of physical factors.

### The regulation of fingering extrusions induces the wound closures independent of the initial curvature

The previously verified linear correlation between the fingering frequency and the wound closure rate is based on the assumption that the initial curvature of the wound is infinity (i.e. straight wounds). Therefore, in order to generalize the relationship between the fingering frequency and the wound closure speeds, consideration of additional effect from the initial curvature values is necessary. As shown in Fig. 4(e), the constituent components of boundary shapes such as *R_initial_*, fingering amplitude (*h*), and fingering frequency (*nf* = *1/FFD*) determine the local curvature of the valley (1/*r_v_’*) that reflects the contractile force (T/*r_v_’*) for wound closures. The mathematical relationship between these 4 variables can be expressed as follows (Fig. S6).

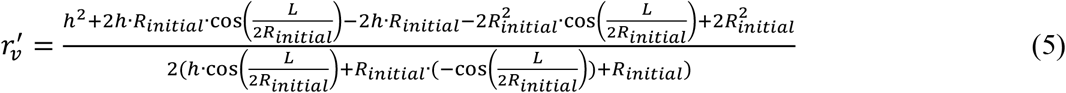

By fitting the data of *R_initial_* and *FFD* to Eq. 5, we obtain the values of coefficient terms, *h* (58.27 μm) and *r_v_’* (89.32 μm), which best estimate the measured values (Fig. S6). The *h* value is already confirmed as an independent value to the *R_initial_* changes (Fig. S1), which ranges from 40 to 70μm, supporting the validity of the optimized value. After fixing the *h* to the optimized value, we can plot the contour map of *r_v_’* according to the *L* and *R_initial_*, followed by laying over the experimental data on the contour map for comparison (Fig. 4(f)). The data points are well distributed on the suggested decaying contour with optimized *h* and *r_v_’* (marked in gray), which confirms that the regulation of finger extrusion by *R_initial_* must cause the constant *r_v_’* to be independent of *R_initial_*. It is notable that the experimental data for *r_v_’* values in the wounds of various *κ_initial_*, indeed, exhibit nonsignificant differences for various initial radii and the value similar to the expected *r_v_’* (marked in gray) (Fig. 4(g)). This result confirms the independent wound closure speeds shown in Eq. (2). These intriguing results imply that the retardance of fingering extrusions in the wounds of larger *ν_initial_* correlates with the conservation of *r_v_’* and closure rate in circular wounds of varying sizes with varying curvatures.

Based on these results, the proposed wound closure steps in Fig. 1(h) can be explained as an orchestration of cell density-driven pressure, cell-cell repulsions, and initial boundary curvature. As shown in Fig. 4(h), the degree of roughness in the wound boundary is influenced by the initial cellular density and is further perturbed by cell division-induced fingering. The cellular density acts as an upstream cue for controlling fingering frequency, and those fingers link the density to the wound closure rate via shaping the curvature of valleys(1/*r_v_’*) Fig. 4(i). Contrary to expectations, the convergence of cells towards the center of the wound caused by negative initial curvature does not result in higher fingering frequency; rather, the frequency is inversely correlated with initial curvature (1/ *R_initial_*). Consequently, this counterbalanced decrease in fingering frequency equalizes the wound closure rate by forming similar local curvatures along the valley independent to the initial curvatures.

### Curvature-driven monolayer structure regulated the fingering extrusions

Finally, we investigate the mechanism for suppressing the fingering extrusions at the higher *κ_initial_* condition. At first, we check the changes in the density-related characteristics while varying the *κ_initial_*. The cellular density near the boundary, that are considered a dominant factor for inducing fluxes and initial fingers, exhibits no significant differences for the different values of *κ_initial_* (Fig. S7(a, b)). Spatiotemporal changes of densities also do not exhibit any distinctive patterns amongst various initial radii (or *κ_initial_*) (Fig. S7(c-e)). When the alignment of traction force, the proposed factor for determining the fingering extrusion region (Fig. S4), is examined, the vector field of traction forces near the boundary is considerably aligned to the radial direction independent of the wound diameter at the initial time as shown in Fig. 5(a). When traction alignments at 0hr and 3 hrs are compared in the wounds of 3 different sizes, the vectors in only the smallest wound (*R_initial_*=150μm) show a noticeable loss in the alignment and random distribution of vectors at 3hr. For quantification of the radial alignment of traction force, the fraction of vectors between −30° and 30° to the radial axis is calculated (Fig. 5(b)). As shown in Fig. 5(c), the proportion of aligned traction vectors has a similar value at the initial time point, but after 3 hrs, the proportion of aligned vectors of the *Rinitiai*=150μm condition significantly decreases. From these results, we propose that the initial development of fingers must occur irrespective of the *κ_initial_* but additional finger formation is retarded by disoriented traction vectors for small wounds of higher negative curvature. Timely maturation of the actomyosin structures whose function is to counterbalance the fingering extrusions^14^, could also contribute to the suppression of finger formations during the closure (Fig. 5(d, e)). Interestingly, after 4 hrs, the radius of multicellular actomyosin curvatures (*R_actomyosin_*) are mostly distributed below 200μm, which coincides with the critical *R_initial_* where the fingering frequency is dramatically decreased (Fig. 5(f)). Based on these data, we postulate that the suppressed finger formation at the higher negative curvature levels may be induced by the maturation of actomyosin cables. Alternatively, from the correspondence between the previously proposed *r_v_’* (Fig. 4 (h)) and the *R_actomyosin_*, the “self-control” of *r_v_’* value may be the consequence of the curvatures of actomyosin rings that induce contraction force for cell migration.

**Fig. 5.**
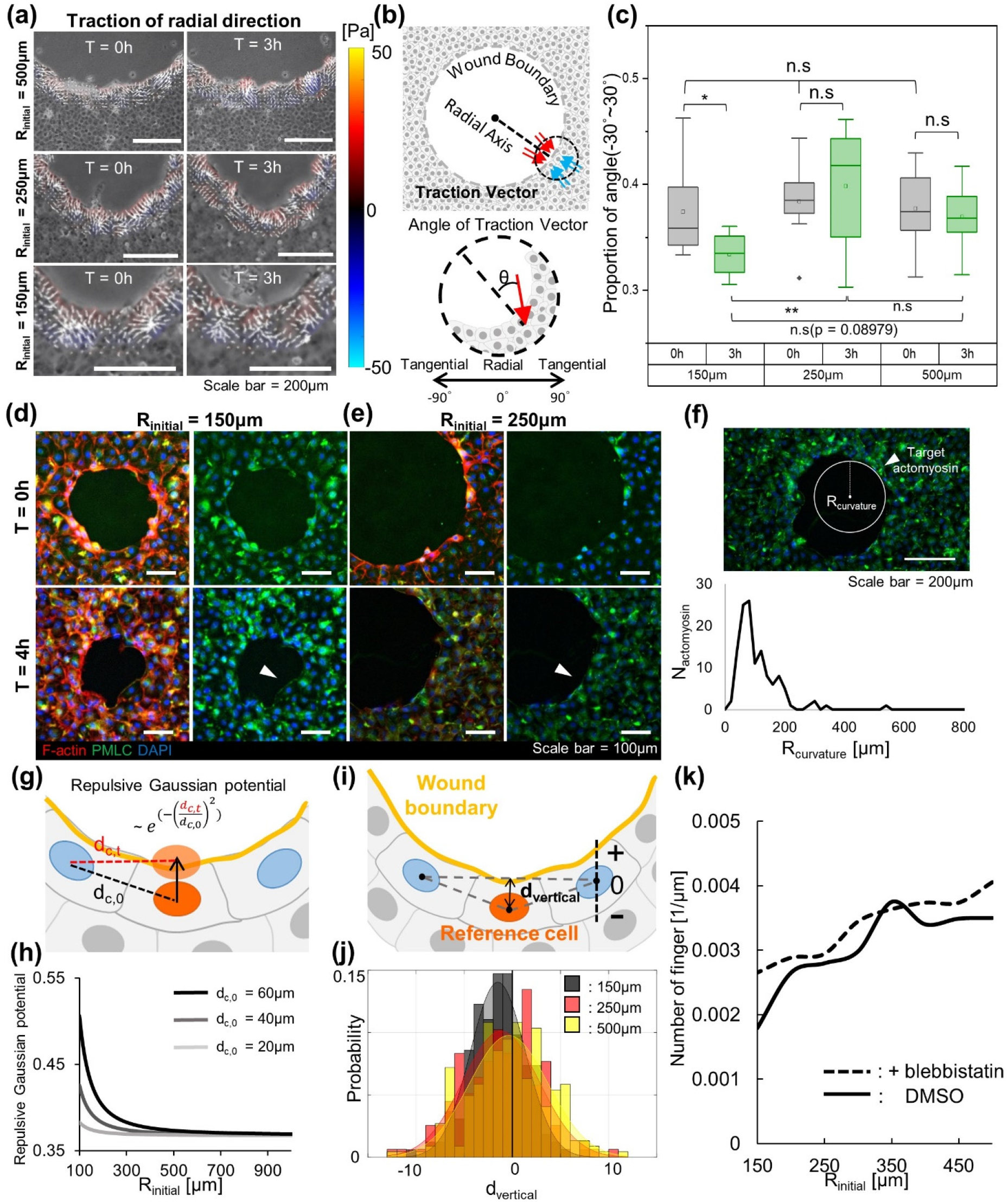
Mechanisms for retarding the fingering extrusions in the wounds with higher initial curvatures. (a) Radial traction force distribution before fingers were formed in circular wounds with various diameters. (b) Definition of the traction angle with respect to the radial directions in the circular wounds. (c) Proportion of traction angles that aligned to the radial directions at 0 and 3 hrs later starting wound closures. (d, e) Immunofluorescence images for the cytoskeletal structures at 0hr and 4hr after closure, which related to the formation of actomyosin rings where the R_initial_ = 150, 250μm, (red = phalloidin, green = myosin II light chain, blue = DAPI), (f) Populations of multicellular actomyosin rings according to the neighboring radius of curvature(R_curvature_), (g) Schematic for determining the repulsive gaussian potential at the wound boundary, (h) Simulation results about the repulsive gaussian potential in terms of initial radius of wounds in various cell densities. (i) Definition of reference cell and distance from the connecting line, the negative sign meant that reference cell is under pressure to behind by adjacent cells. (j) Histogram of the distance from the connecting line according to the wound diameter. (k) Changes in the number of fingers per unit length according to the initial radius when the blebbistatin (25μm) was treated or not (DMSO: n = 3, samples: 84, blebbistatin: n =2, samples: 52).

The position of cells can also have the suppressing effect for finger development. In various cellular migration models, the intercellular distance has been considered as an important factor that contribute in determining the intercellular energy between neighboring cells. One type of energy from the cell-cell interactions is repulsive potential with Gaussian shapes (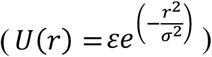), that captures the mildly repulsive system like cells^46^. In the initially curved wound boundary, the boundary cells would increase this repulsive potential as they move forward due to the intrinsic curvature effect (Fig. 5(g)). As shown in Fig. 5(h), the mathematical model predicts a dramatic increase in the repulsive potential at the smaller *R_initial_* condition. When the cell-cell distance is decreased to *d_c,0_* = 20μm, which represents the extremely high density of cells, the effect of the increase in potential becomes insignificant when *R_initial_* = 150μm. As the cells are known to generate the repulsion force by neighboring cells close enough^34,35^, the relative position of a cell with respect to the other neighbors can determine the direction of the total repulsion force, allowing us to predict the direction of cell protrusion (Fig. 5(i)). As shown in Fig. 5(j), the wounds of all three sizes show higher distributions of backward-directed repulsions force vectors that work against the extrusion of fingers.

To confirm the importance of the multicellular actomyosin ring and repulsion potentials in suppressing finger formation, we conducted an analysis while inhibiting myosin activity with blebbistatin (25μM). As shown in Fig. 5(k), the average number of fingers was higher in the blebbistatin-treated condition, confirming the significant suppressive function of actomyosin rings in finger extrusion. However, both conditions showed a dramatic decrease in fingering frequency for smaller wounds with a closer cell-cell distance, indicating that repulsive potential based on cell position also plays a critical role in modifying finger extrusion. In these cytoskeletal-based analyses, we acknowledge that the physical forces due to negative curvature make it challenging for boundary cells to form fingers.

## Discussion

In this study, we explored the spontaneous emergence of finger-valley structures that appear along the boundary during the closure of an epithelial monolayer. During the wound closure, the fingers are formed by leader cells that crawl faster than neighboring cells, and the cells in the concave valley regions between fingers accelerate to catch up by utilizing contractile forces to smoothen the roughness. If the speed disparities between the leaders in the fingers and neighboring cells were to be maintained, the finger amplitudes would continue to grow, resulting in increasing boundary roughness. The roughness index (RI) value rose quickly as fingers emerge, saturated at a constant level while fingers and valleys move at a similar level, and fell until fingers were suppressed and the wound was closed by the contraction of the actomyosin cable. We identified the driving factors for fingering extrusions as cellular density and the global curvature of the initial boundary, demonstrating how the finger-valley structure immediately affected wound closure rates via inter-controlled crawling and boundary contraction by external factors like initial curvatures and cellular density.

In order to investigate the impact of fingering characteristics on wound closure rates, we developed a simple mathematical model for the curvature along the fingers. Our findings revealed that the correlation between fingering frequency and wound closure rate was stronger than the relationship between cellular density and wound closure rate, which is typically the primary mechanism for collective cell migration. We also found that cellular density had a stronger correlation with fingering frequency, indicating that cellular density plays an upstream role in finger generation. We confirmed that the gradient of cellular density induced diverging cell fluxes, which were then transmitted to the extruding location of fingers via cell-cell adhesions. Cell division also played a crucial role in increasing fingering frequency by forming divergence and reducing the velocity correlation length in the monolayer (Fig. S8). Based on previous work by Vishwakarma M. et al., which explains the occurrence of additional fingers when the mechanical stress correlation lengths are shorter, we hypothesized that the dividing rate or initial cell density would determine the physical correlation lengths for velocity or mechanical stress between cells that ultimately determine the occurrence of fingers^33^.

The initial concave curve at the wound boundary induced the converging cell flux from the monolayer, predicting more fingering extrusions as the curvature rose. However, fingering extrusion was suppressed at higher curvatures, which aligned with a mathematical relationship for uniform curvature values, regardless of the initial curvatures. Additionally, experiments revealed insignificant differences in local valley curvatures between samples with different initial global curvature values. These results demonstrate that the fingering frequency must be finely regulated to meet a certain valley curvature value irrespective of the global boundary shapes. In addition, the wound boundary maintained a similar closing speed regardless of the initial curvature of the wound, employing a “self-control” mechanism likely to maximize closure efficiency by regulating energy costs. By controlling the finger protrusion and valley contraction to maintain the overall roughness of the boundary at a moderate wound closure speed, undesired stretching of cells and rupture of fingers can also be prevented. Along the boundary, the multicellular actomyosin arc formed, acting as obstacles against protruding lamellipodia to counterbalance the crawling fingers. We have also observed a similarity in length scale between the characteristic radius of valley curvature and the radius of the actomyosin arc, suggesting the possible existence of a characteristic length of the actomyosin arc that induces contractile migrations.

Furthermore, our research has shown that the fingering extrusion is determined by the relative positions of the cells, which reflect the physical potential of each constituent cell. This phenomenon is similar to the change in melting temperature at solid-liquid interfaces, which is influenced by exterior curvatures as described by the Gibbs-Thomson equations^36^. We also have observed that the protrusion of fingers driven by cellular flux can be explained by adjusting the particle movements with a soft repulsion force. This provides further insight into the mechanisms behind fingering extrusion and highlights the importance of considering the physical properties and interactions of individual cells in these processes. Overall, our approach to the complex biological event is significant because it can be generalized to broader physical phenomenon. Therefore, the proposed mechanisms are anticipated to be utilized in various contexts to simplify the complex biophysical systems that feature rough boundaries with multiple protrusions.

## Material and methods

### Cell Culture

MDCK (Mardin Darby canine kidney) cell line, derived from normal epithelial cells of the dog kidney, was cultured in DMEM (Gibco, USA) media supplemented with 10% fetal bovine serum and 1% penicillin/streptomycin. In both culture and live imaging systems, cells were maintained in a humified incubator with 5% CO2 and 37°C.

### Fabrication and adaptation of silicone stencils for making wounds

The PDMS (polydimethylsiloxane) silicone oil (Sylgard 184, Downing corning) was utilized to make the stencils with various embossed shapes for wounds. The PDMS elastomer was cured in a 10:1 ratio with a curing agent and poured into the SU-8 wafer fabricated by the photo-lithography methods (MIcroFIT, Seongnam, Korea). After removing the residual gases inside of the PDMS through the Vacuum container, we cured the PDMS for at least 3 hrs in the oven (>65°C). The cured PDMS stencil was gently detached from the wafer and cut into the block that contains a similar shape of embossed structures with various sizes. Each block was treated with the oxygen plasma to make a hydrophilic surface for preventing the bubble generation from loads of cell suspensions. The PDMS blocks were attached to glass-bottom dishes, and the cell suspension was gently put between the blocks and glasses. Then, we incubated cells until they form the monolayer except for the embossed structures.

### Live cell imaging

Phase-contrast images for observing the cell movements and fluorescent images for displacements of the substrate were taken every 10~15 minutes using a 5X objective lens. All experiments on the live-cell imaging were conducted on the Axiovert 200M (Carl Zeiss) microscope with an incubating system (37°C and 5% CO2).

### Quantification of curvature along the wound boundary

Local curvatures of valley regions were measured from a circle passing three points: the center and both ends of valleys. The curvature value was a reciprocal number of the circle radius, and the center of the circle determined the sign of curvature. If the center point is located at the exterior of the cell layer (wounded region), the curvature sign would be negative.

### Visualization of cell motilities via the DPIV method

The quantification of cell migration was conducted by visualizing cellular motilities. For the visualization of cells, we adjusted the DPIV (digital particle image velocimetry) method to the sequence of phase images. The commercial software, Image Velocimetry Tool for MATLAB, calculated the cross-correlation between cell images^37^. The double-pass PIV by the Fast Fourier Transform (FFT) window deformation algorithm was used for the calculation. We determined the first window size for 32·32 pixels (64·64μm2) and the second window size of 16·16 pixels (32·32μm2) with an 8 pixels (16 μm) interval that is similar to the length of single cells. In this way, the velocity vector fields of the cell monolayer were gained. Other motility factors, like divergence and correlation length, could be calculated from the vector data.

### Proliferation controls of cell

To control the proliferation of cells, we adapted the thymidine, DNA synthesis inhibitor, treatment. Six hrs before starting the wound closure experiment, thymidine was added to the cell group to a final concentration of 2mM for waiting; the cells were arrested at the G1/S boundary.

### Immunostaining of cell cytoskeletons

For the immunofluorescence staining, 4% paraformaldehyde was used to fix the cell layer for 20 min at room temperature, then the cells were rinsed with PBS at least three times. The fixed cell layer was permeabilized with 0.5% Triton X-100 in PBS for 15min with an ice pack and washed three times with PBS. To block the unspecific binding, we incubated the cell layer in the 3% BSA in PBS in the incubator for 30mins. After blocking, the cells were incubated in the primary antibody in 3% BSA with a 1:100 ratio at 4°C overnight. Next, the samples were rinsed three times with 3% BSA and additionally incubated with a secondary antibody (Alexa Fluor 488 goat anti-rabbit antibody (A11008, Thermo Fisher Scientific)) in 3% BSA with a 1:200 ratio at room temperature for 2 hrs. For staining the actin cytoskeletons and nucleus, Rhodamine-phalloidin (R415, Thermo Fisher Scientific) in 3% BSA with a 1:50 ratio and 4’,6-diamidino-2-phenylindole (DAPI, Thermo Fisher Scientific) were employed, respectively.

### Hydrogel-based traction force visualization substratum

Mapping forces of cells require the soft gel that can be deformed by the cellular traction force. The deformable soft gel was made from the polyacrylamide (PA) gel, which could regulate its stiffness by changing the composition. For capturing the deformation of gels, the fluorescent beads layer between cells and PA gel was adopted. The mixture of PA gel and fluorescent beads (diameter = 0.5um) was polymerized on the glass bottom dish and simultaneously centrifuged to pull the beads to the top surface of the gel. Next, the polymerized PA gel was treated with sulfosuccinimidyl-6-(4-azido-2-nitrophenylamino) hexanoate (Sulfo-SANPAH; Proteochem) 1 mg/ml in 50 mM HEPES buffer (Life Technologies) and added 50 μg/ml collagen type I (PureCol; Advanced BioMatrix) for being an adaptable gel for cell living.

### Traction force microscopy via Fourier transformation

The map for the traction force fields of the cell monolayer was measured from the deformation of PA-gels, which is visualized by the displacement of the fluorescent beads layer. We used the unconstrained Fourier transform traction microscopy by utilizing the previously published algorithms^38–40^ adapted in MATLAB software.

## Statistical Analysis

Statistical data were analyzed using Origin and Excel software. The box plotting of data was conducted by using the Origin software, and the Excels and MATLAB software handled other plots. For the statistical results, the data were compared using the Mann-Whitney test and One-Way ANOVA in Origin software. Statistical significance is marked as * P < 0.05, ** P < 0.01, *** P < 0.001.

## Acknowledgments

J.J. Fredberg’s lab generously provided TFM and MSM codes at Harvard T.H. Chan School of Public Health. This research was supported by National Research Funding granted by the Korean Government (NRF-2017R1A2B2007673, NRF-2020M3A9E4039658) and by the KAIST (Basic Science Research Program for faculty members).

## AUTHOR INFORMATION

### Notes

The authors declare no competing financial interest.

