## Supplementary material for "The fingering patterns in the epithelial layer control the gap closure rate via curvature-mediated force": Supplemental Figure.pdf

### FIG. S1

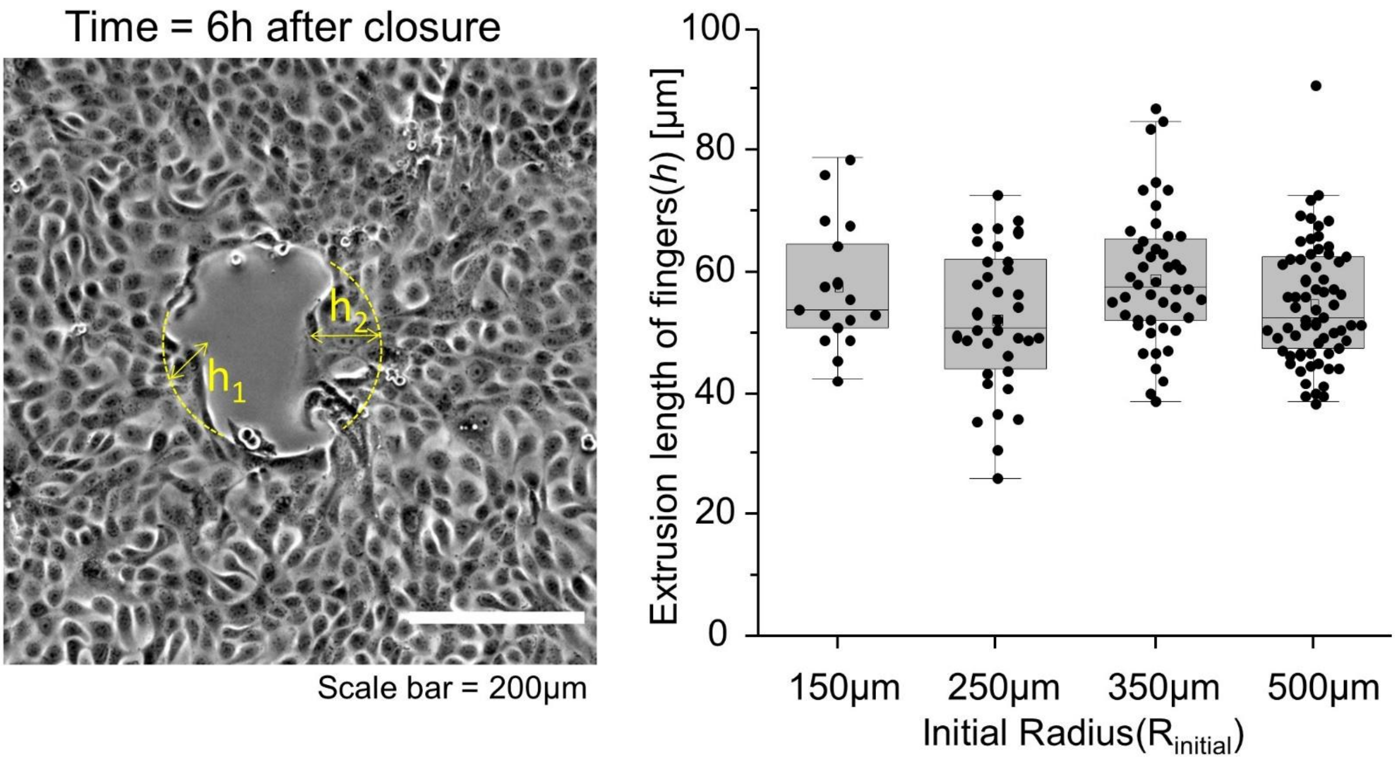


**Figure S1.** The measurement of extrusion length of fingers after 6 hours from the start of closing, the results showed less significant differences according to the initial radius of wounds.

### FIG. S2

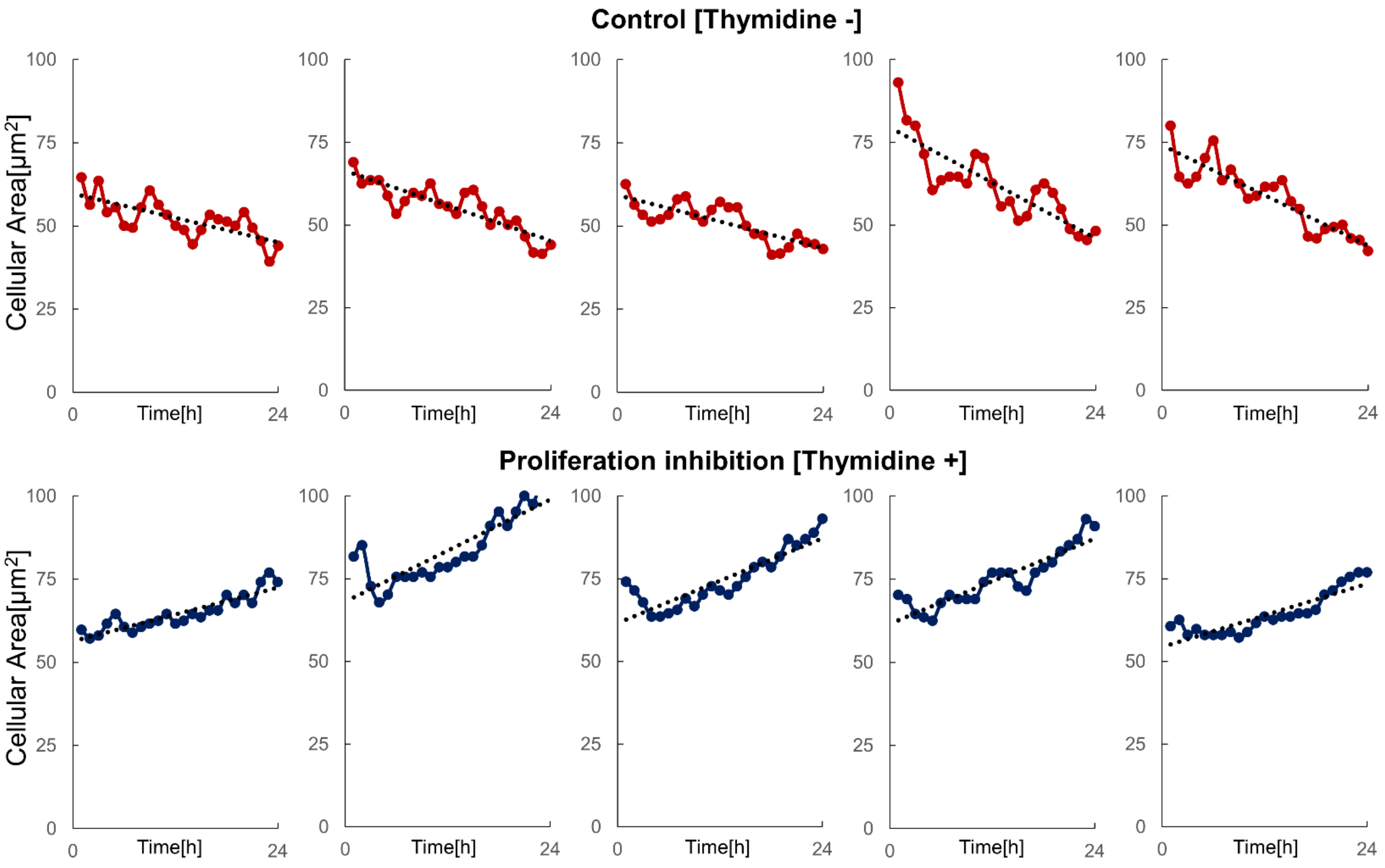


**Figure S2.** The results of the temporal fluctuations of mean area cells in 200um from the boundary, when the thymidine is treated or not.

### FIG. S3

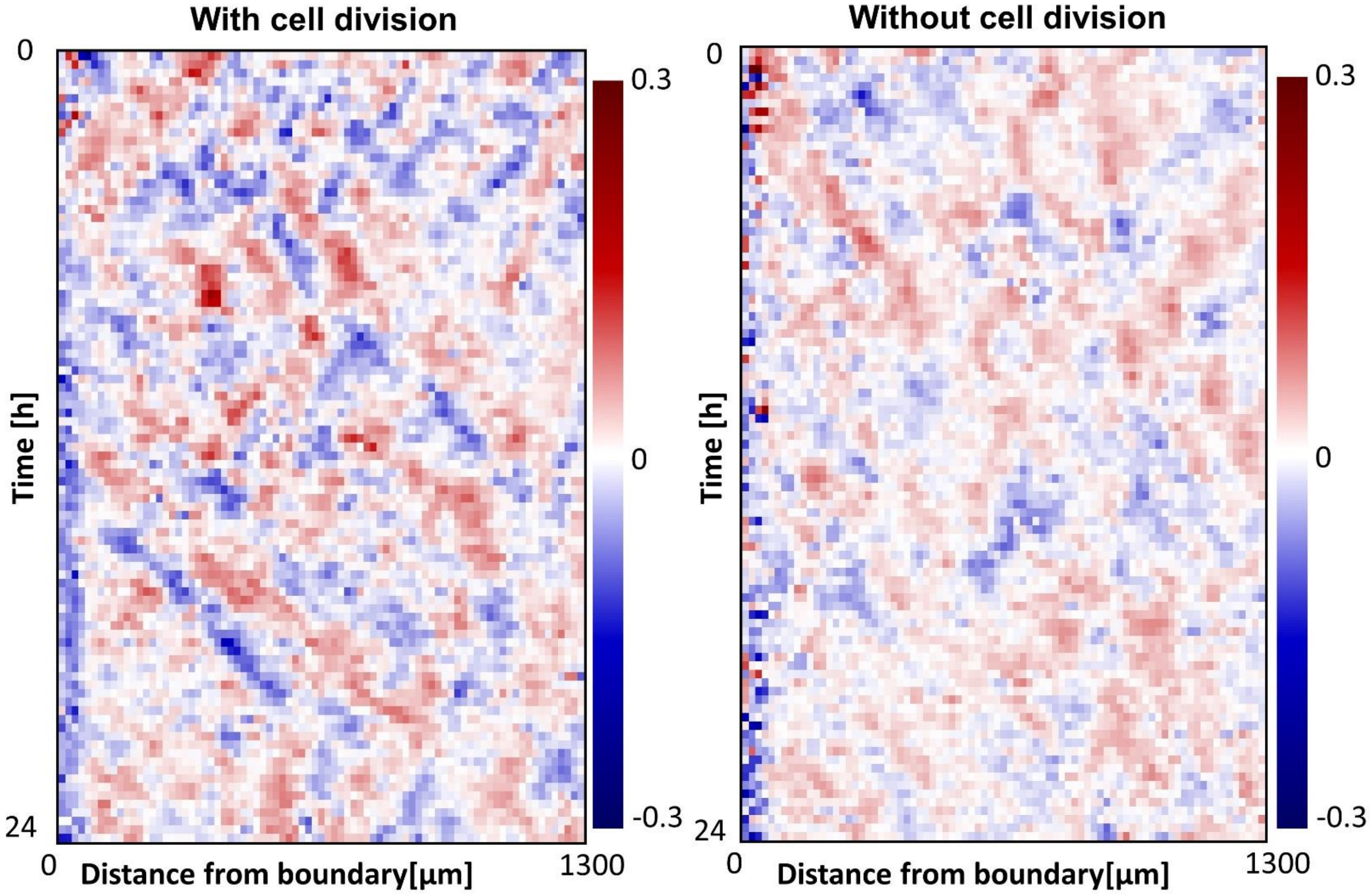


**Figure S3.** Spatiotemporal changes of divergence maps according to the cell division conditions.

### FIG. S4

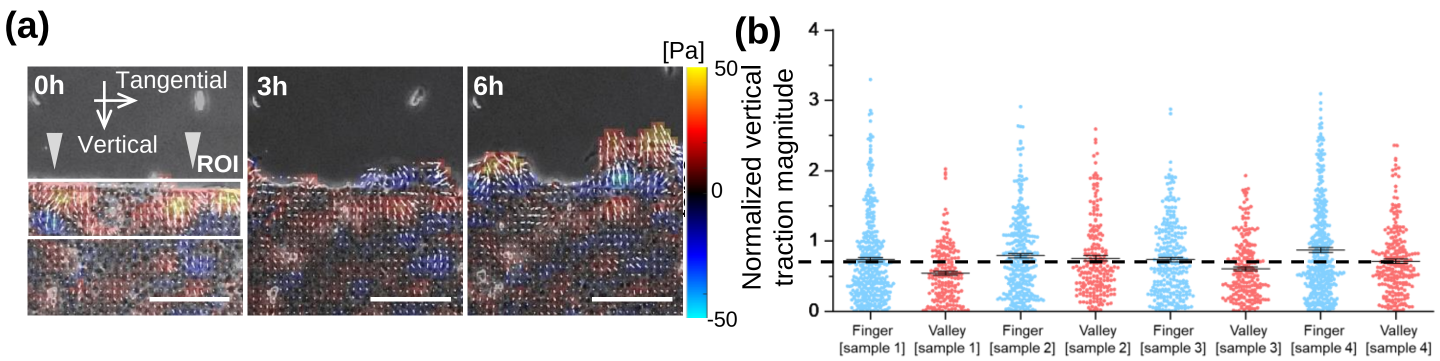


**Figure S4. Distribution of traction vectors near the fingering extrusion site.** (a) Development of fingering structures in relation to the traction distribution. Gray triangles indicate the fingering extrusion sites and rectangular is ROI for measuring the average traction magnitude. (b) Distribution of normalized vertical traction magnitudes within the 100µm behind the finger(blue) and valley(pink) region.

### FIG. S5

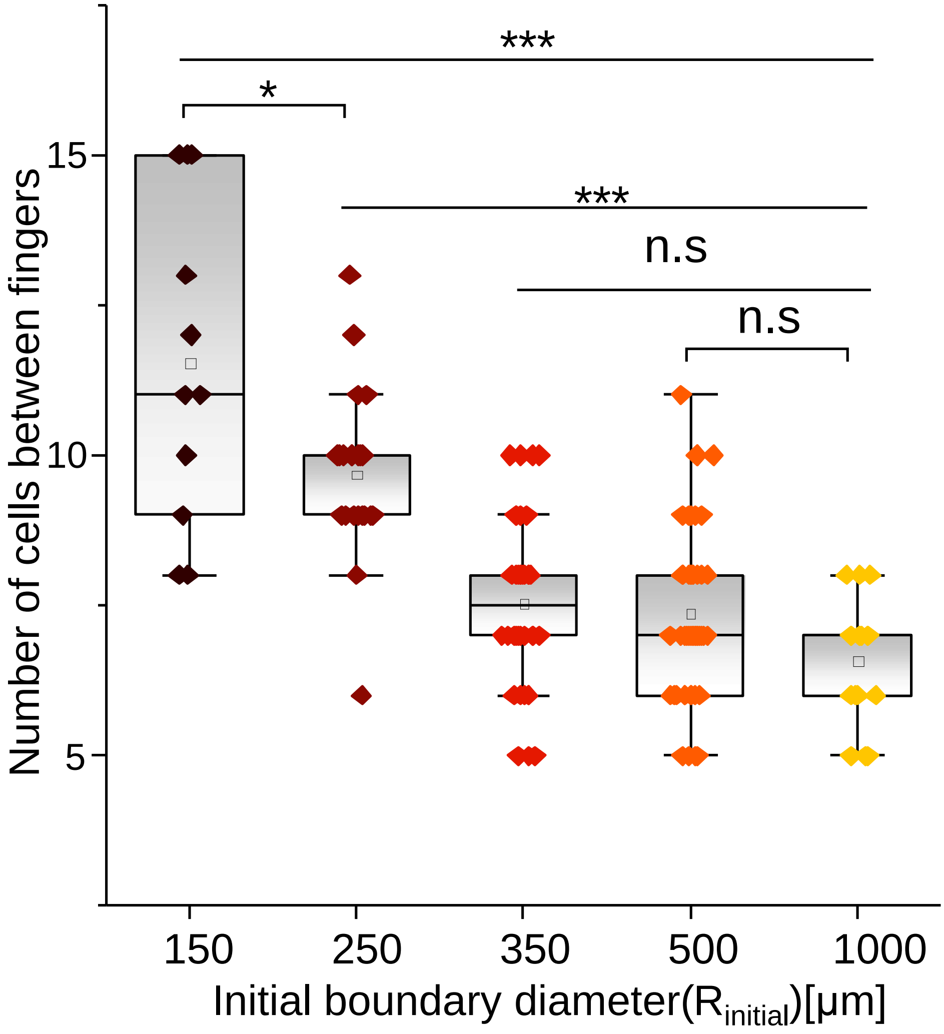


**Figure S5.** Number of cells between fingers according to the initial diameter of wounds

### FIG. S6

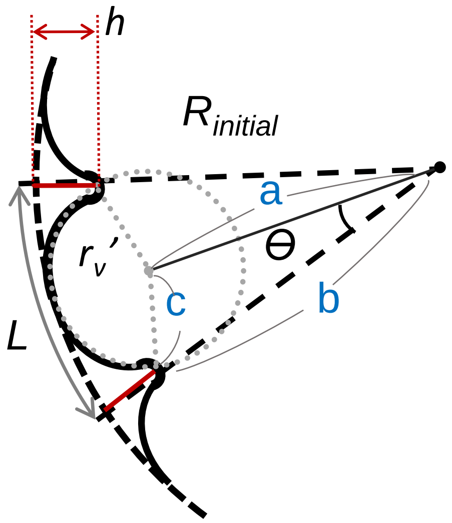


$$\frac{1}{2} a \cdot b \cdot sin\theta = \sqrt{p \cdot (p-a) \cdot (p-b) \cdot (p-c)}$$

$$when, p = \frac{1}{2}(a+b+c)$$

$$a = R_{initial} - r_v'$$

$$b = R_{initial} - h$$

$$c = r_v'$$

$$\theta = \frac{L}{2R}$$

$$r_v{}' = \frac{h^2 + 2hRcos\left(\frac{L}{2R}\right) - 2hR - 2R^2\cos\left(\frac{L}{2R}\right) + 2R^2}{2\left(h\cos\left(\frac{L}{2R}\right) + R\left(-\cos\left(\frac{L}{2R}\right)\right) + R\right)}$$

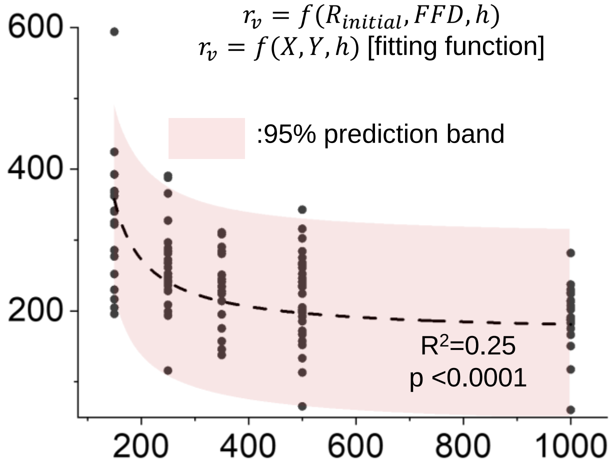


Fitted value

| $h$ | $r_v$ |
|---|---|
| 58.27 μm | 89.32 μm |

**Figure S6.** Formulation of equation for the valley curvatures(after fingers were formed) of valleys when cell boundary is initially curved and the fitted results with experimental data.

### FIG. S7

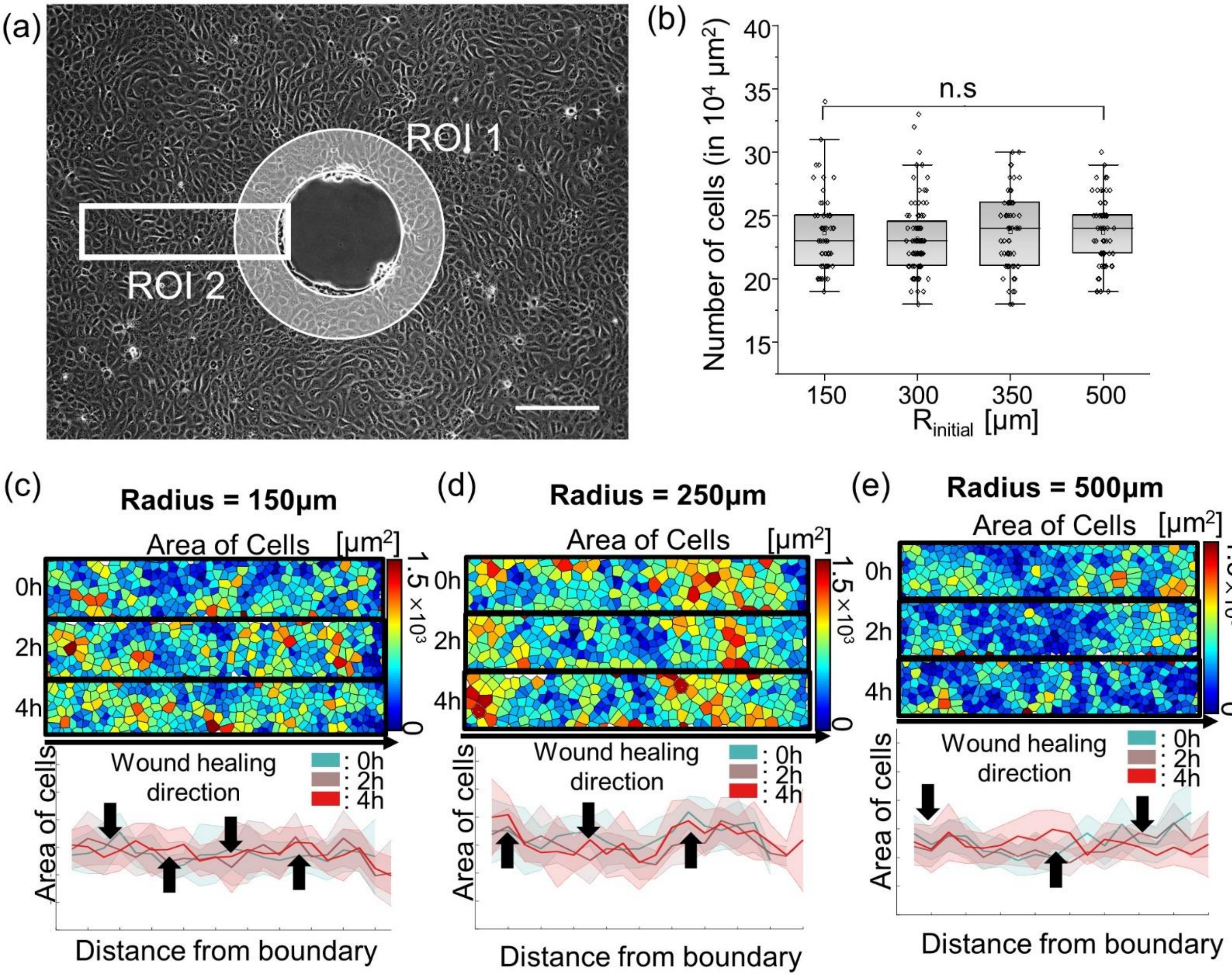


**Figure S7 Density properties according to the curvature changes of circular wounds.** (a) Location of region of interests for analyzing the density properties of cells. (white circle : region of interest for measuring the cellular density near the boundary, yellow rectangular : region of interest for analyzing the perturbation of cell morphologies), (b) Initial density of cells according to the diameters of circular wounds. The initial density did not show significances amongst the conditions. (c~e) Morphology distribution of cells according to the wound diameters, the shapes of cells were assumed by the Voronoi tessellation algorithms for the center position of cells.

### FIG. S8

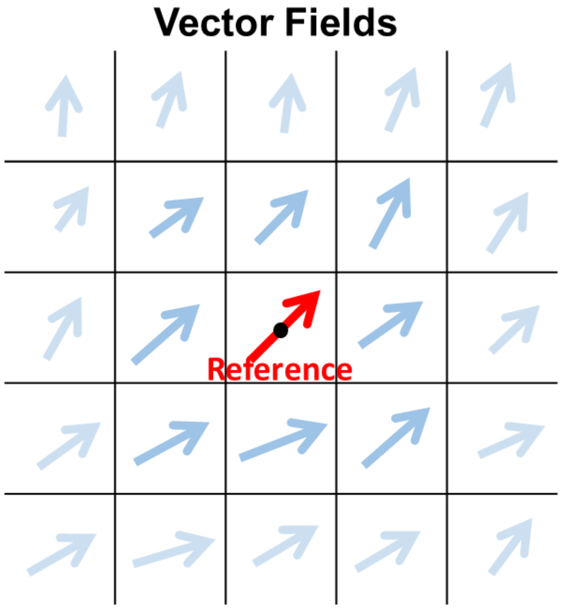


***Correlation value***

$$= \frac{1}{N}\sum_{i=1}^{N}\frac{u_{reference}\cdot u_i + v_{reference}\cdot v_i}{\left(\sqrt{u_{reference}^2 + v_{reference}^2}\right)\cdot\left(\sqrt{u_i^2 + v_i^2}\right)}$$

Where, $N$ is the total number of grid with same distance from the reference grid, and $i$ indicates the each grid.

***Correlation length***

$$= Distance\ from\ the\ reference\ grid,\quad where\ \boldsymbol{Correlation\ value} > 0.9$$

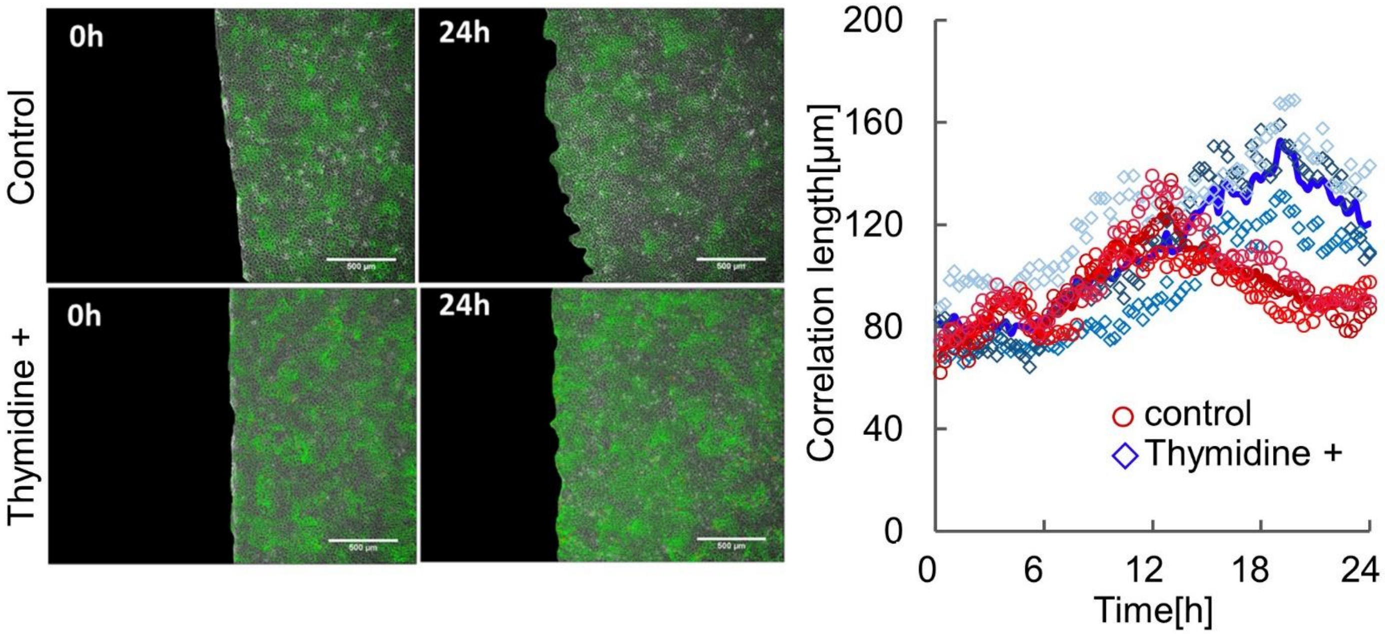


**Figure S8.** Results for the correlation length of the cell velocity quantified by the PIV when the thymidine is treated or not
